# Assessing the additional effects of piperonyl butoxide (PBO) exposure on *Anopheles mosquitoes*

**DOI:** 10.64898/2026.06.15.732125

**Authors:** Rosheen Mthawanji, Polius Gerazi Pinda, Steven Gowelo, Themba Mzilahowa, Hilary Ranson, Christopher M. Jones

## Abstract

Insecticide treated nets (ITNs) remain one the frontline tool in malaria control in Africa. The synergist piperonyl butoxide (PBO) has been incorporated into ITNs to combat pyrethroid resistance and these pyrethroid-PBO ITNs are now in widespread use across Africa. PBO is known to inhibit the degradation of pyrethroids and other insecticide classes, but little is known about other biological or behavioural effects of PBO exposure. This work assessed the impact of PBO on *Anopheles gambiae* s.l. male longevity as well as female host-seeking and blood feeding behaviours using laboratory assays. PBO had an immediate impact on blood feeding of female *An. gambiae* s.l. whereas the effect was not observed 24h post-exposure. Kaplan-Meier survival curves showed a significant difference in mosquito longevity post-exposure to PBO with male mosquitoes living nearly three times as long if exposed to control papers compared to PBO. In olfactometer assays, after exposure to control papers, control mosquitoes were significantly more attracted to host-odours, however, this attraction preference was reversed following PBO exposure. This study shows that in experimental assays, PBO has other effects on mosquito physiology and this is likely to contribute to the impact of pyrethroid - PBO ITNs in vector control programmes.

## Introduction

In the last 15 years the routine widespread distribution of ITNs across malaria endemic areas has contributed to the prevention of over 450 million cases [1]. All ITNs in use contain pyrethroid insecticides [2]. Unfortunately, the high coverage of ITNs along with indoor residual spraying (IRS), and use of pyrethroids in agricultural and household products, has led to the evolution of pyrethroid resistance [3]. Consequently, the efficacy of ITNs has been undermined and this has led to an urgent call for new tools with different modes of action in malaria vector control [4]to avoid the reversal of epidemiological gains from ITN programmes[5].

In response to growing pyrethroid resistance, manufacturers have produced ITNs containing the insecticide synergist piperonyl butoxide (PBO). PBO acts by inhibiting the enzyme group, cytochrome P450s which detoxify or sequester insecticides within insects [7]. The role of P450s in pyrethroid resistant mosquitoes is well established across malaria vector species [8][9]. The addition of PBO alongside a pyrethroid increases the killing effect on malaria vectors that express P450-based resistance mechanisms [11]. Multiple experimental hut trials have shown an increased efficacy of pyrethroid-PBO ITNs against resistant mosquitoes compared to pyrethroid-only nets [12,13,14,15]. In addition to insecticide resistance, P450s play a wide variety of metabolic roles within insects including hormone synthesis and protection from other xenobiotics [16][17. Due to the broad function of P450s it is likely that exposure to this compound may impact other aspects of mosquito physiology and through direct or indirect mechanisms [9].Any indirect effects that reduce the ability of mosquitoes to transmit malaria may enhance the overall effectiveness of PBO ITNs [18][19].

Here, our aim was to examine the effects of PBO in the absence of a pyrethroid, on male longevity, female blood-feeding, and host-seeking of *An. gambiae* s.l. using laboratory assays.

## Methods

### Study locations and mosquito strains

The study was conducted in two locations: the Malaria Alert Centre (MAC) laboratory in Blantyre (southern Malawi) and the Ifakara Health Institute (IHI) in Tanzania.

Experiments conducted at MAC used the susceptible *An. gambiae s*.*s* (Kisumu strain). Kisumu strain reared under standard insectary conditions humidity. In Ifakara, susceptible insectary reared *An. arabiensis* was used which was reared under standard insectary conditions.

### Exposure to PBO in WHO tube assays

In the MAC laboratory, *Anopheles gambiae* s.s. (Kisumu) were used to determine the effects of PBO alone using the WHO tube bioassay [20]. This was carried out using 4% PBO impregnated papers (made at LSTM) with untreated papers used as the control. Batches of 20 to 25 3-to-5-day old mosquitoes [29] were exposed for one hour as part of three different objectives.

i. *The effect of PBO on blood-feeding one day post exposure.* 3-5-day old males and females (placed in separate tubes) were exposed to either control orPBO papers for one hour and then placed in cages and allowed to mate for 24 hours at a 1:2 male:female ratio. Following the mating period, female mosquitoes were exposed to an artificial blood feeding system (Hemotek) for 45 to 60 minutes where the mosquitoes were fed cow blood. The number of mosquitoes that fed (F) and did not feed (NF) was recorded.
ii. *The effect of PBO on blood-feeding immediately after exposure.* 3-5-day old female mosquitoes were held in the WHO tubes for one hour and exposed to PBO or control papers. Immediately after exposure female mosquitoes were blood-fed as above. The number of mosquitoes that fed (F) and did not feed (NF) was recorded.
iii. *Effect of PBO on male mosquito longevity.* A total of 100 1-3-day old male mosquitoes were exposed to 4 % PBO for one hour in the WHO tubes in replicates of 25 individuals. Post-exposure mosquitoes were placed into separate falcon tubes with 10% sugar water. Knockdown andmortality were measured every 24 hours.

### Olfactometer bioassay

The response of mosquitoes to human odour after exposure to PBO was investigated using a Y-tube olfactometer at [21]. The olfactometer consists of a stem and two treatment chambers where odours are located (Figure 1B). In this experiment 3-5-day old lab reared female *An. arabiensis* mosquitoes were starved for 5 hours prior to 1h exposure with PBO or control papers using WHO resistance bioassay tubes. Immediately after exposure, mosquitoes were aspirated into the stem of an olfactometer horizontally placed on a lab bench under insectary conditions. Each chamber contained either odour cues (worn socks) or negative control (empty chamber). Once the socks were placed on the end, mosquitoes were released into the chambers for 5 minutes and the preference for the treatment chamber or control was timed and recorded. After 5 minutes, the experiment was stopped, all the gates were closed and mosquitoes in each section were counted, this experiment was done in 5 replicates.

**Figure 1.**
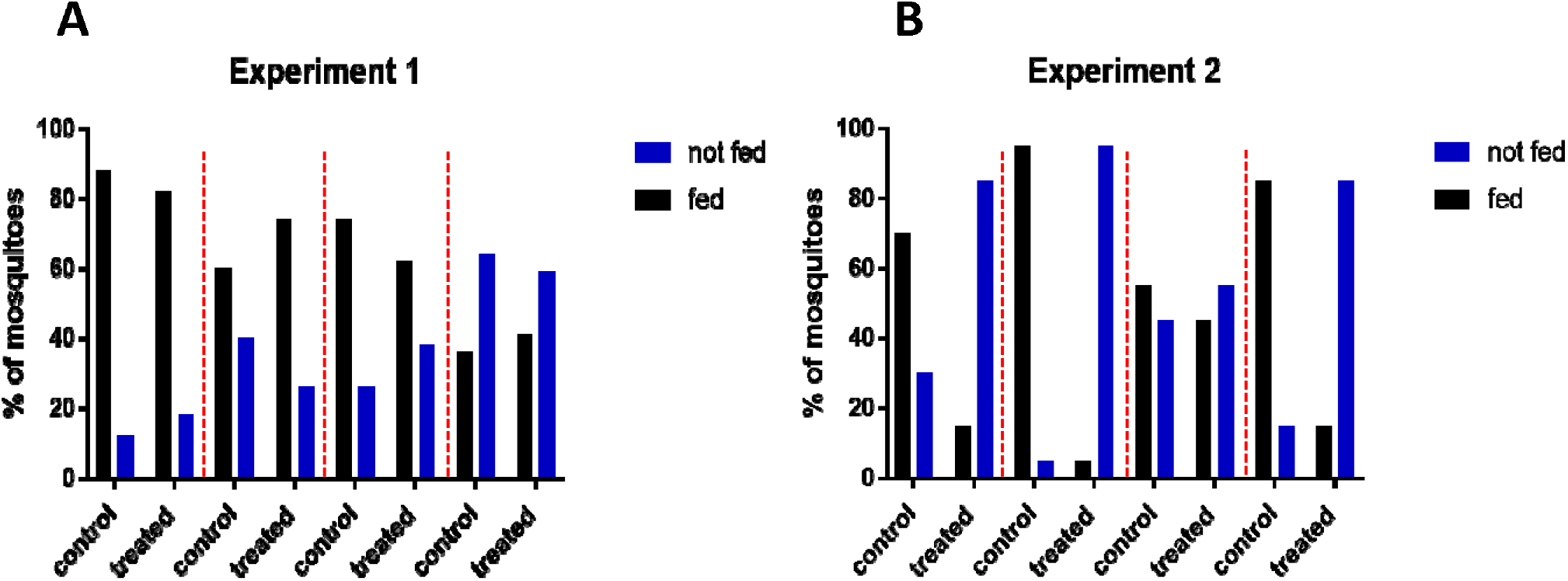
The blood feeding response of *An. gambiae* Kisumu post exposure to PBO papers. (A) Results for experiment 1 show the percentage of female mosquitoes blood-feeding 24 h post exposure to PBO. (B) Experiment 2 shows the percentage of mosquitoes blood feeding immediately after exposure to PBO. F = blood-fed. NF = non blood-fed females. Separate replicated experiments are shown by dotted red lines.

## Data analysis

Survival data were analysed using Kaplan Meier curves and Gehan-Breslow-Wilcoxon test and the log-rank test within GraphPad prism [22]. The Y-tube olfactometer data were analysed using an exact binomial test under the null hypothesis that mosquitoes would enter either the treatment or control chambers in a 50:50 ratio. The olfactometer analyses were performed using R Statistical Software (Version 4.0.2) [23].

## Results

## Figures

### Blood feeding of female *An. gambiae* and longevity of males after exposure to PBO in the absence of a pyrethroid

Three experiments were performed using the WHO tube assays to analyse the impact of PBO (in the absence of a pyrethroid) on blood feeding and male longevity. Mosquitoes exposed to either PBO or control papers and allowed to feed 24 h later, showed no difference throughout the replicates in blood-feeding following exposure to either non-treated or PBO papers (χ^2^=6.04, df=7, P = 0.54) (Figure 1A). By contrast, where blood feeding was allowed immediately after exposure there was a significant reduction in feeding following PBO exposure (χ^2^=429.0, df=7, P < 0.001) (Figure 1B). Kaplan-Meier survival curves showed a significant difference in mosquito longevity post exposure to control and PBO papers with male mosquitoes living nearly three times as long if exposed to control papers rather than those treated with PBO (χ ^2^ = 66.1, df = 1, p < 0.001) (Figure 2).

**Figure 2.**
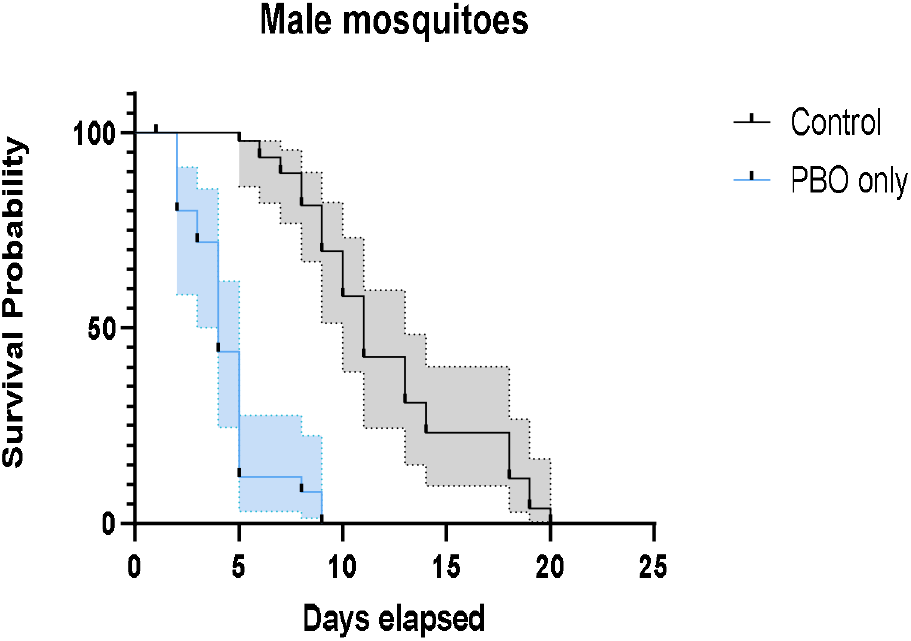
Kaplan-Meier curves in response to exposure to PBO for male *An. gambiae* s.l. graphs. Probability of survival is shown as a function of age post exposure.

### Attraction to host-odours following PBO exposure

Host-seeking by laboratory reared female *An. arabiensis* was disrupted following a 1h exposure to PBO papers in olfactometer bioassays. After exposure to control papers, responding mosquitoes were significantly more attracted to the host-odour chamber (73.3%, 95% CI = 61.9-82.9%, P < 0.001), however, this attraction preference was reversed following PBO exposure (10.2%, 95% CI = 3.4-22.2%, P < 0.001) (Figure 3). It is noteworthy that a much higher percentage of mosquitoes (N = 51%) did not respond at all once aspirated into the olfactometer after PBO exposure compared with the control (25%).

**Figure 3.**
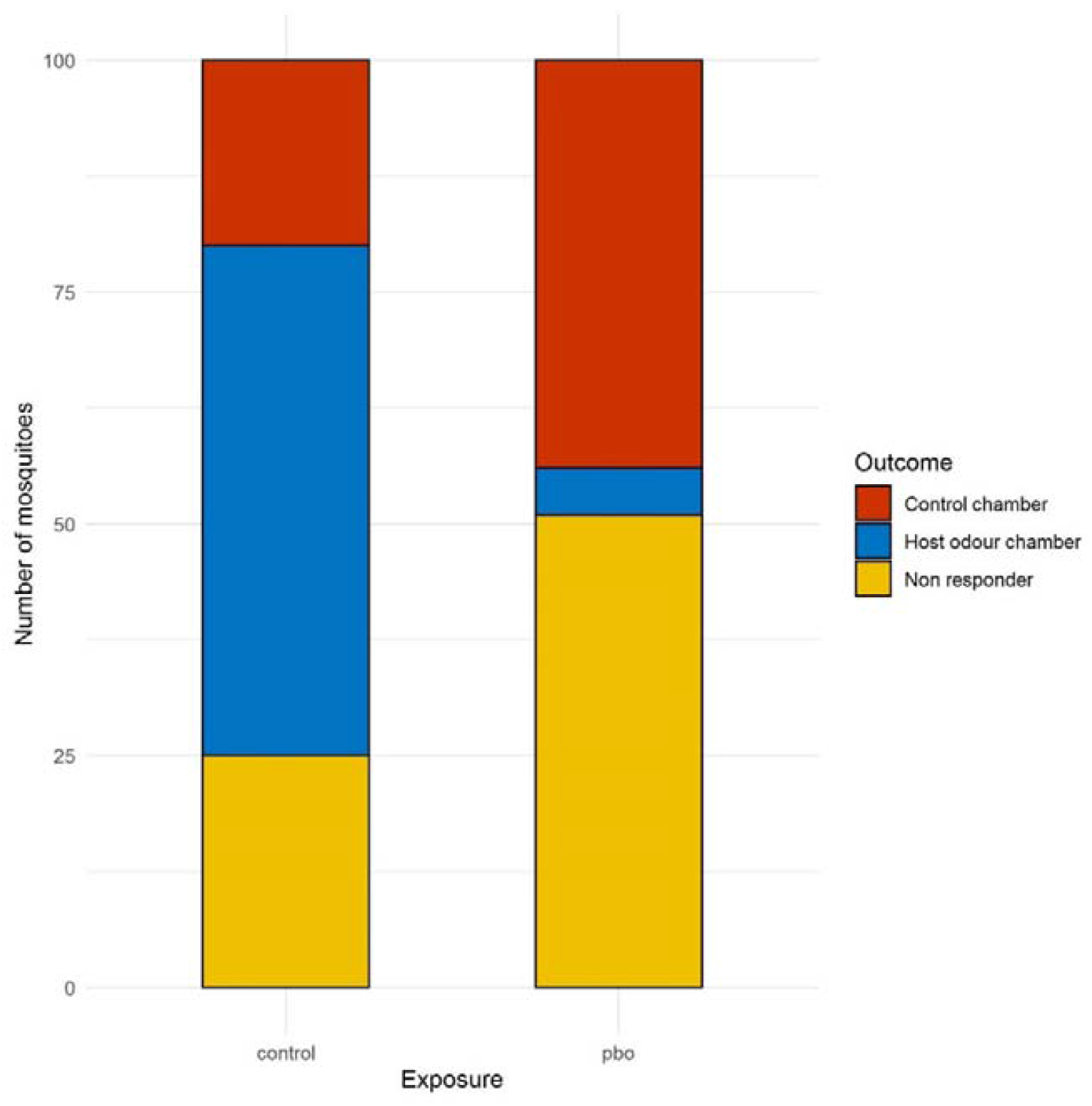
Response of laboratory reared *An. arabiensis* females in an olfactometer following exposure to PBO or control WHO papers. Following aspiration at one end of the stem of the olfactometer, mosquitoes were given a choice between host odours (socks) or a control chamber. Non-responders were removed from the data analysis and a total 100 mosquitoes were exposed to each treatment in batches of 25.

## Discussion

A systematic review of experimental hut and clinical trials concluded that pyrethroid-PBO ITNs increase mosquito mortality and reduce blood feeding rates in areas of high pyrethroid resistance [24]. In Tanzania and Uganda, two independent cluster randomised trials demonstrated that PBO ITNs reduce malaria parasite prevalence compared with standard ITNs [25]. In 2021, approximately half of all ITNs procured for distribution in Africa were pyrethroid-PBO ITNs [26] Maintaining the efficacy of PBO-ITNs is therefore essential, especially now national control programmes are distributing PBO-ITNs as part of routine mass campaigns. The main aim of this study was to determine the immediate and sub-lethal impacts of exposure to PBO in the absence of a pyrethroid against *An. gambiae* s.l. mosquitoes. The results provide some further evidence that the incorporation of PBO onto ITNs imparts additional fitness costs against *An. gambiae* s.l. mosquitoes.

PBO is a synergist inhibiting the cytochrome p450 enzyme group of which over 100 are described in the major mosquito vectors [26]. Even in the absence of an insecticide it is plausible that exposure to PBO could have an additive negative impact on mosquito physiology or behaviour given the broad range of functions p450s play in insect physiology. Exposure of female susceptible *An. gambiae* to PBO for one hour had a significant impact on the ability of mosquitoes to blood-feed. This effect was only observed immediately after exposure and was absent 24h later (Fig. 1). Similar studies [27][28] exhibit mosquitoes spending less time feeding on the synergist net than on ITNs without the synergist as well as untreated nets, showing detrimental effects on blood feeding. Similar complementary observations on the impact of PBO exposure on host-seeking behaviour were observed in the olfactometer assays (Fig. 3) with increased proportions of mosquitoes either failing to respond to host-odours or not responding at all.

Finally, our experimental work showed that PBO exposure reduces male mosquito survival. The majority of exposure to PBO under field conditions will, of course, be borne by female mosquitoes but given the increasingly widespread distribution and use of PBO-nets, and evidence for indoor observations of male *An. gambiae* (e.g. indoor swarming) [28], it seems likely that at least a small proportion of the male mosquito population will be exposed to this compound. Perhaps more interestingly, is the stark decline in survival following PBO exposure suggesting that there is at least some indirect physiological impact on male longevity although this could be exaggerated by the fragility of male mosquitoes under experimental conditions.

## Conclusion

Overall, we present evidence that PBO in the absence of a pyrethroid (or other insecticide) has an impact on aspects of mosquito physiology. This laboratory-based study demonstrates that the impact of PBO-ITNs probably goes beyond insecticide-induced mosquito mortality and factors such as impaired host-seeking, reduced blood-feeding and reduced male survival post-contact may contribute to the effectiveness of PBO-ITNs in addition to their obvious toxicity against resistant mosquito malaria vectors.

## Acknowledgements

This work was funded in whole, by the Wellcome Trust International Masters Fellowship award to Rosheen Mthawanji [208300/Z/17/Z].

## Author contributions

RM performed the experiments. RM, CMJ, TM and HR conceived the ideas and designed the experiments. RM and CJ analysed the data. RM wrote the first draft of the paper. All authors contributed to the reviewing of the drafts and gave approval to publish results

## Ethical approval

The study ‘Assessing the effects of Piperonyl Butoxide (PBO) exposure on malaria vector fitness in Chikwawa district, Malawi (ATEPBO)’ was approved by the College of Medicine research and ethics committee (COMREC), permit number P.07/19/2738

## Consent for publication

All authors have approved the submitted manuscript

## Open access

For the purpose of open access, the author has applied a creative commons by public copyright licence to any author accepted manuscript version arising from the submission

## Data availability

The data generated during the study are available. **{WILL NEED TO DEPOSIT DATA IN REPOSITROY E.G. DRYAD]**

## References

1. World Health Organisation (2019) World Malaria Report 2019. WHO. Geneva.

2. WHO. Global technical strategy for malaria 2016–2030. Geneva: World Health Organization; 2015.

3. Bhatt, S., Weiss, D., Cameron, E., Bisanzio, D., Mappin, B., Dalrymple, U., Battle, K., Moyes, C., Henry, A., Eckhoff, P., Wenger, E., Briët, O., Penny, M., Smith, T., Bennett, A., Yukich, J., Eisele, T., Griffin, J., Fergus, C., Lynch, M., Lindgren, F., Cohen, J., Murray, C., Smith, D., Hay, S., Cibulskis, R. and Gething, P. (2015). The effect of malaria control on Plasmodium falciparum in Africa between 2000 and 2015. Nature, 526(7572), pp.207–211.

4. World Health Organization. (2017). Report of the twelfth WHOPES working group meeting, WHO/HQ, Geneva, 8-11 December 2008: review of bioflash GR, permanet 2.0, permanet 3.0, permanet 2.5, lambda-cyhalothrin LN. World Health Organization.

5. Tungu, P., Magesa, S., Maxwell, C., Malima, R., Masue, D., Sudi, W., Myamba, J., Pigeon, O. and Rowland, M. (2010). Evaluation of PermaNet 3.0 a deltamethrinPBO combination net against Anopheles gambiae and pyrethroid resistant Culex quinquefasciatus mosquitoes: an experimental hut trial in Tanzania. Malaria Journal, 9(1), p.21.

6. Jones, G. (1998). Piperonyl Butoxide. Burlington: Elsevier.

7. Feyereisen, R. (2011). Arthropod CYPomes illustrate the tempo and mode in P450 evolution. Biochimica et Biophysica Acta (BBA) - Proteins and Proteomics, 1814(1), pp.19–28.

8. Churcher TS, Lissenden N, Griffin JT, Worrall E, Ranson H: The impact of pyrethroid resistance of the efficacy and effectiveness of bed-nets for malaria control in Africa. Elife 2016, 5.

9. Stevenson BJ, Pignatelli P, Nikou D, Paine MJI. 2012. Pinpointing P450s associated with pyrethroid metabolism in the dengue vector, Aedes aegypti: developing new tools to combat insecticide resistance. PLoS Negl. Trop. Dis. 6, e1595. 10.1371/Journal.pntd.0001595 (doi:10.1371/Journal.pntd.0001595). [PMC free article] [PubMed] [CrossRef] [Google Scholar]

10. Report of the eleventh WHOPES working group meeting [Internet]. Geneva: World Health Organization; (2008). Report of the eleventh WHOPES working group meeting [Internet]. Geneva: World Health Organization;. [online] Available at: http://www.who.int/iris/handle/10665/69732 [Accessed 12 Aug. 2018].

11. Viana, M., Hughes, A., Matthiopoulos, J., Ranson, H. and Ferguson, H. (2016). Delayed mortality effects cut the malaria transmission potential of insecticideresistant mosquitoes. Proceedings of the National Academy of Sciences, 113(32), pp.8975–8980

12. Akoton, R., Tchigossou, G., Djègbè, I., Yessoufou, A., Atoyebi, M., Tossou, E., Zeukeng, F., Boko, P., Irving, H., Adéoti, R., Riveron, J., Wondji, C., Moutairou, K. and Djouaka, R. (2018). Experimental huts trial of the efficacy of pyrethroids/piperonyl butoxide (PBO) net treatments for controlling multi-resistant populations of Anopheles funestus s.s. in Kpomè, Southern Benin. Wellcome Open Research, 3, p.71.

13. Koudou, B., Koffi, A., Malone, D. and Hemingway, J. (2011). Efficacy of PermaNet® 2.0 and PermaNet® 3.0 against insecticide-resistant Anopheles gambiae in experimental huts in Côte d’Ivoire. Malaria Journal, 10(1), p.172.

14. Protopopoff, N., Mosha, J., Lukole, E., Charlwood, J., Wright, A., Mwalimu, C., Manjurano, A., Mosha, F., Kisinza, W., Kleinschmidt, I. and Rowland, M. (2018). Effectiveness of a long-lasting piperonyl butoxide-treated insecticidal net and indoor residual spray interventions, separately and together, against malaria transmitted by pyrethroid-resistant mosquitoes: a cluster, randomised controlled, two-by-two factorial design trial. The Lancet, 391(10130), pp.1577–1588

15. Corbel, V., Chabi, J., Dabiré, R., Etang, J., Nwane, P., Pigeon, O., Akogbeto, M. and Hougard, J. (2010). Field efficacy of a new mosaic long-lasting mosquito net (PermaNet® 3.0) against pyrethroid-resistant malaria vectors: a multi-centre study in Western and Central Africa. Malaria Journal, 9(1), p.113

16. Ranson, H., N’Guessan, R., Lines, J., Moiroux, N., Nkuni, Z. and Corbel, V. (2011). Pyrethroid resistance in African anopheline mosquitoes: what are the implications for malaria control?. Trends in Parasitology, 27(2), pp.91–98

17. Allossogbe, M., Gnanguenon, V., Yovogan, B., Akinro, B., Anagonou, R., Agossa, F., Houtoukpe, A., Padonou, G. and Akogbeto, M. (2017). WHO cone bio-assays of classical and new-generation long-lasting insecticidal nets call for innovative insecticides targeting the knock down resistance mechanism in Benin. Malaria Journal, 16(1).

18. Parker, J. E. A., Angarita-Jaimes, N., Abe, M., Towers, C. E., Towers, D., & McCall, P. J. (2015). Infrared video tracking of Anopheles gambiae at insecticide-treated bed nets reveals rapid decisive impact after brief localised net contact. Scientific Reports, 5(March), 1–14. 10.1038/srep13392

19. Parker, J. E. A., Angarita Jaimes, N. C., Gleave, K., Mashauri, F., Abe, M., Martine, J., … McCall, P. J. (2017). Host-seeking activity of a Tanzanian population of Anopheles arabiensis at an insecticide treated bed net. Malaria Journal, 16(1). 10.1186/s12936-017-1909-6

20. Daimon T, Kozaki T, Niwa R, Kobayashi I, Furuta K, Namiki T, Uchino K, Banno Y, Katsuma S, Tamura T et al: Precocious Metamorphosis in the Juvenile Hormone-Deficient Mutant of the Silkworm, Bombyx mori. Plos Genetics 2012, 8(3)

21. WHO. Test procedures for insecticide resistance monitoring in malaria vector mosquitoes. Geneva: World Health Organization; 2013

22. Meza, F.C., Roberts, J.M., Sobhy, I.S. et al. Behavioural and Electrophysiological Responses of Female Anopheles gambiae Mosquitoes to Volatiles from a Mango Bait. J Chem Ecol 46, 387–396 (2020). 10.1007/s10886-020-01172-8

23. Prism version 8.0.0 for Windows, GraphPad Software, San Diego, California USA, http://www.graphpad.com

24. R Core Team. (2013). R: a language and environment for statistical computing. R foundation for statistical computing. Vienna, Austria URL http://www.R-project.org.

25. David J, Ismail HM, Chandor-Proust A, & Paine MJ. Role of cytochrome P450s in insecticide resistance: impact on the control of mosquito-borne diseases and use of insecticides on EarthPhil. Trans. R. Soc. B3682012042920120429 10.1098/rstb.2012.0429

26. https://allianceformalariaprevention.com/working-groups/net-mapping/

27. Akoton, R., Tchigossou, G., Djègbè, I., Yessoufou, A., Atoyebi, M., Tossou, E., Zeukeng, F., Boko, P., Irving, H., Adéoti, R., Riveron, J., Wondji, C., Moutairou, K. and Djouaka, R. (2018). Experimental huts trial of the efficacy of pyrethroids/piperonyl butoxide (PBO) net treatments for controlling multiresistant populations of Anopheles funestus s.s. in Kpomè, Southern Benin. Wellcome Open Research, 3, p.71.

28. Barreaux P, Koella JC, N’Guessan R, Thomas MB. Use of novel lab assays to examine the effect of pyrethroid-treated bed nets on blood-feeding success and longevity of highly insecticide-resistant Anopheles gambiae s.l. mosquitoes. Parasit Vectors. 2022 Mar 28;15(1):111. doi: 10.1186/s13071-022-05220-y. PMID: 35346334; PMCID: PMC8962112.

29. Gleave K, Lissenden N, Richardson M, Choi L, Ranson H. Piperonyl butoxide (PBO) combined with pyrethroids in insecticide-treated nets to prevent malaria in Africa. Cochrane Database Syst Rev. 2018;1101:2776.

30. Barreaux P, Koella JC, N’Guessan R, Thomas MB. Use of novel lab assays to examine the effect of pyrethroid-treated bed nets on blood-feeding success and longevity of highly insecticide-resistant-Anopheles gambiae s.l. mosquitoes. Parasit Vectors. 2022 Mar 28:15(1);111. Doi:10.1186/s13071-022-05220-y. PMID;pudmed:35346334: PMCID; pudcentral: PMC8962112

31. David, J., Ismail, H., Chandor-Proust, A., & Paine, M. (2013). Role of cytochrome P450s in insecticide resistance: impact on the control of mosquito-borne diseases and use of insecticides on Earth. Philosophical Transactions Of The Royal Society B: Biological Sciences, 368(1612), 20120429. 10.1098/rstb.2012.0429

